# Phylogeography of *Lanius senator* reveals conflicts between alpha taxonomy, subspecies ranges and genetics

**DOI:** 10.1101/2021.10.27.466041

**Authors:** Martina Nasuelli, Luca Ilahiane, Giovanni Boano, Marco Cucco, Andrea Galimberti, Marco Pavia, Emiliano Pioltelli, Arya Shafaeipour, Gary Voelker, Irene Pellegrino

## Abstract

Implementing the effort in understanding biogeographic distribution patterns and taxonomic limits within animal groups is crucial for addressing several challenges of modern zoology. Although avian phylogeography has been deeply investigated within Western Palearctic, several families, such as shrikes, still display complicated or neglected biogeographic patterns both between and within species, thus requiring further investigations. The Woodchat Shrike (*Lanius senator*) is a long-distance migratory species that exhibits three morphologically well-recognizable subspecies, whose boundaries have never been molecularly investigated. Here, we aimed to define the phylogeographic structure of *Lanius senator* throughout its breeding range and assess the genetic coherence with the phenotypically described subspecies. We assembled a collection of 34 samples mainly from breeding populations of each subspecies and analyzed them at four mtDNA and two nuDNA markers. We did not find a clear phylogenetic structure with nuclear Ornithine Decarboxylase (ODC) and myoglobin intron 2 (MYO), while all the four mtDNA loci (i.e., ND2, COI, *cytb* and Control Region) highlighted two main haplogroups, one including both the nominate subspecies *L. s. senator* and *L. s. badius* and the second consistent with *L. s. niloticus* only from the easternmost part of the range. Surprisingly, individuals phenotypically assigned to *L. s. niloticus* from Israel were genetically assigned to the *senator/badius* haplogroup. Moreover, genetic distances showed intermediate values between inter-intraspecies diversity usually found in Passerines. We estimated a divergence time among the two haplogroups around 800 kya (549 - 1.259 kya HPD). Our findings showed a mismatch in subspecies assignment using morphology and genetic information and a marked differentiation between the eastern *L*.*s. niloticus* and all the other *L. senator* populations.

## Introduction

Understanding taxonomic boundaries by assessing genetic structure and biogeographic distributions of natural populations is essential to promoting biodiversity conservation (e.g., Lohman et al. 2010; Huntley et al. 2019; Galimberti et al. 2021), and to include phylogenetic inference and speciation dynamics in modern zoology (Degnan and Rosenberg 2009; Shi and Rabosky 2015). In birds, a growing body of literature suggests that many recognized subspecies, which were named via often subtle variations in plumage or morphology, are either poorly circumscribed in terms of geographic distribution, or are invalid when assessed via molecular methods (e.g., White et al. 2013; Wojczulanis-Jakubas et al. 2014; Semenov et al. 2017).

For example, the Western Palearctic (WP) is one of the most investigated biogeographic regions in terms of avian phylogeography, where to date 145 bird species have been the focus of molecular analyses (Pârâu and Wink 2021). Despite fairly good coverage in terms of species analyzed (i.e., approximately 20% of the 720 WP breeding species; Pârâu and Wink 2021), the WP continues to present challenges related to 1) species-complexes that have been poorly (and perhaps never) investigated, and 2) additions from unsampled areas to previous studies (e.g., Ilahiane et al. 2021). The second challenge is particularly important in relation to the southern areas of Europe, several of which are known glacial refugia. It is clear that these refugia played a decisive role in shaping intra and interspecific geographical variations, with phylogeographic studies identifying Iberia, Southern Italy or the Balkans as regions where populations accumulated genetic differences (e.g., Brito 2005; Pellegrino et al. 2015; Drovetski et al. 2018a; Albrecht et al. 2020). Strong intraspecific differentiation is rare in migratory WP species (Pârâu and Wink 2021), being instead largely confined to sedentary or short-distance (intra-Palearctic) species.

Shrikes (Passeriformes; Laniidae) are one group that has received considerable attention through phylogeographic studies, and 3 of the 8 species with a range that includes western palearctic regions have been investigated (Gonzalez et al. 2008; Olsson et al. 2010; Kvist et al. 2011; Padilla et al. 2015; Pârâu et al. 2019). Different phylogenetic studies conducted on *Lanius* species showed complicated biogeographic and evolutionary patterns both between and within species. In some cases, the distinction between species and subspecies remains poorly understood, and genetic studies show unclear and complex relationships among morphologically described taxa (species and subspecies, Peer et al. 2011; Fuchs et al. 2019). One case where possible confusion still exists in terms of subspecies taxonomy is the Woodchat Shrike *Lanius senator*. The Woodchat Shrike is a polytypic species which breeds in the Western Palearctic (Figure 1), with all populations migrating long distance to winter in sub-Saharan Africa (Shirihai and Svensson 2018). Several subspecies have been described, based on breeding, migrating, or wintering individuals. A detailed list of proposed taxa and synonyms over time by several authors is shown in Table S1. The various subspecies designations have been based on subtle plumage differences or on biometric characteristics. As a consequence, most recent taxonomic treatments recognize only three subspecies: *Lanius senator senator, L. s. badius* and *L. s. niloticus* (del Hoyo et al. 2008; Dickinson and Christidis 2014; Shirihai and Svensson 2018; Clements et al. 2021; Gill et al. 2021), while a few exceptions consider *L. s. rutilans* as a valid taxon (Cramp and Perrins 1993; del Hoyo and Collar 2016). None of these taxonomic decisions were based on genetic data.

**Figure 1.**
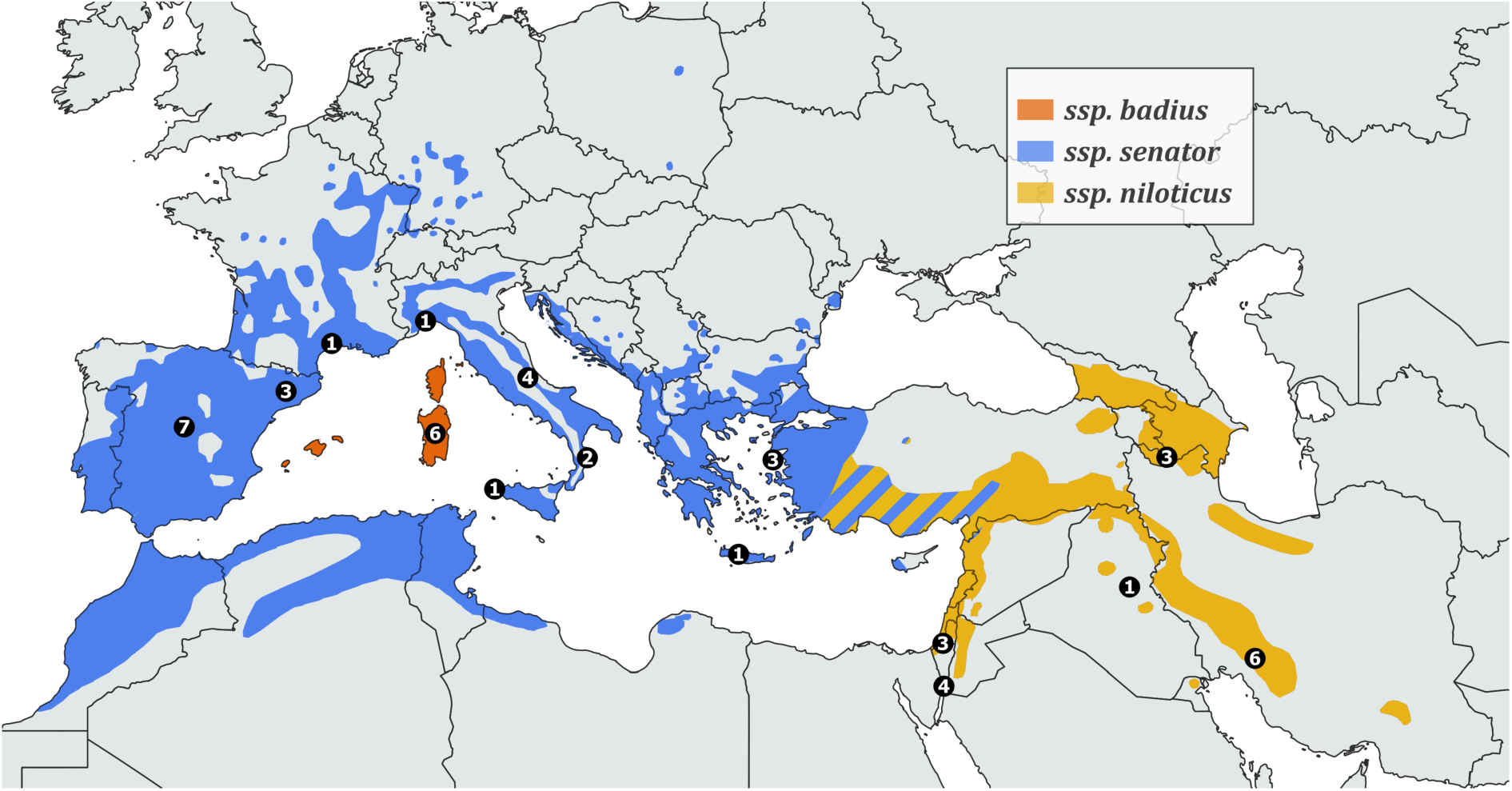
Breeding range of *Lanius senator*. Coloured areas delimit the distribution of subspecies *L. s. senator, L. s. badius, L. s. niloticus*, as shown in the legend. Black dots define the samples size for each locality (specific details on sampling sites are reported in Table 1 & Supplementary Material S2). Wintering range is not shown, although we included a Genbank sequence from Gambia in the analysis. The map was made with QGIS 3.16.10 (https://qgis.org) and it was based on BirdLife International and Handbook of the Birds of the World (2020).

*Lanius senator senator* is widespread across the Mediterranean basin from the Iberian Peninsula to Western Turkey and winters in Sub-Saharan Africa from Senegal to Sudan (Figure 1). It is characterized by a moderately large white primary patch and no or very little white at the base of the tail feathers, a characteristic visible only in hand. Individuals from Eastern Greece and Western Turkey show on average more white on the wings and tail. *L. s. badius* is endemic to the Balearics, Corsica and Sardinia and winters from West Africa east to Cameroon. It differs from the other subspecies in having no (or a tiny) white spot on the primaries and in having on average a darker back without white on tail coverts and feathers. It is also generally larger with a stronger bill and shows less sexual dimorphism than the other subspecies. *L. s. niloticus* breeds from Cyprus, southern and eastern Turkey eastward through the Middle East and Transcaucasia to southeastern Iran, and winters in the extreme south of the Arabian Peninsula, and East Africa. It differs from the other subspecies in having a larger white base on all the tail feathers (up to 32 mm) which is often visible in the field, a larger white patch on primary feathers and whiter underparts giving the impression of a more black-and-white bird than the other subspecies. The taxon *L. s. rutilans*, described from Senegal and reported as breeding in the Iberian Peninsula and North Africa was recognized as valid by comparatively few authors until recently (see Table S1), but it does not show any clear difference in plumage or biometry (Shirihai and Svensson 2018) relative to the three well-accepted taxa (Shirihai and Svensson 2018).

Distributions of above-mentioned taxa in some cases overlap, for example, while Roselaar (1995) confirmed the presence of both *L. s. senator* and *L. s. niloticus* as common breeders in western or southeastern Turkey, respectively, he also reported the presence of a mixture zone or areas where the taxonomic attribution of individuals is uncertain, especially in the Anti-Taurus Mountains (southern and eastern Turkey) and in the coastal area close to the Syrian border (south-central Turkey). In addition, Shirihai (1996) indicated that while *L. s. niloticus* is the only breeding taxon in Israel, where it is also common during migration, *L. s. senator* is also a migrant there, albeit scarce (around the 10% of the total migrating Woodchat Shrikes), and restricted to a narrow migration window (mainly from mid-April to the beginning of May). Finally, the presumed *L. s. rutilans* would have a partially overlapping distribution with *L. s. senator* in the north-east of the Iberian Peninsula.

In this study, using a multi-locus genetic approach (i.e., two nuclear and four mitochondrial markers) we (1) investigated, for the first time, the phylogeographic structure of *Lanius senator* throughout most of the breeding range of the species, and (2) inferred the genetic relationships and differentiation among populations of the three widely accepted subspecies (*L. s. senator, L. s. niloticus* and *L. s. badius)*.

## Materials and Method

### Sample collection

We collected 34 samples of *Lanius senator* from different localities spread across the breeding range of the species (Figure 1). In particular, we sampled *L. senator* in northern and southern Italy including Sardinia, Armenia, Iran and we obtained samples from museum specimens from north-eastern and central Spain (Table 1, Table S1). To increase the sample set we also added a few samples of migrating individuals collected on Marettimo Island (near Sicily) and Eilat (Southern Israel). These samples were included into the analysis since their taxon attribution in the field was certain, being based mainly on plumage features, following Svensson (1992). The biological samples consisted of muscle tissue (N=21) from museum collections, blood (N=10) collected by us in the field, and DNA extracts (N=3) (see Supplementary Information 1 for sampling details). Muscle tissues were preserved in 96-100% ethanol, while blood was preserved in ethanol or Queen’s Lysis buffer (Seutin et al. 1991). In both cases samples were stored at -20°C until laboratory analysis.

**Table 1.** List of specimens investigated in this study with voucher/accession number, subspecies, sex, sampling locality and origin. MZB: Museu de Ciències Naturals de Barcelona; MCC: Museum of Naturali History of Carmagnola; W.I.N.E.S.: collection from TAMU/UNIUPO/MusNatHist Carmagnola; NHMO: Natural History Museum University of Oslo; MNCN: Museo Nacional de Ciencias Naturales Madrid

| Voucher specimen/accession number | Subspecies | Sex | Country | Locality | Museums |
| --- | --- | --- | --- | --- | --- |
| IT 0813 | *Lanius senator badius* | F | Italy | Sardinia, Telti | W.I.N.E.S. |
| IT 0820 | *Lanius senator badius* | M | Italy | Sardinia, Telti | W.I.N.E.S. |
| IT 1090 | *Lanius senator senator* | M | Italy | Calabria, Caccuri | W.I.N.E.S. |
| IT 1092 | *Lanius senator senator* | M | Italy | Calabria, Caccuri | W.I.N.E.S. |
| IT2867 | *Lanius senator senator* | M | Italy | Abruzzo, Volpe | W.I.N.E.S. |
| IT2868 | *Lanius senator senator* | M | Italy | Abruzzo, Volpe | W.I.N.E.S. |
| MNCN6732 | *Lanius senator senator* | F | Spain | Madrid, Alcalá de Henares | MNCN |
| MNCN10812 | *Lanius senator senator* | M | Spain | Ciudad Real, Pueblonuevo del Bullaque | MNCN |
| woodchat 1 | *Lanius senator niloticus* | M | Iran | Kohgiluyeh and Boyer-Ahmad Province, Yasuj | Yasouj University |
| woodchat 2 | *Lanius senator niloticus* | F | Iran | Kohgiluyeh and Boyer-Ahmad Province, Yasuj | Yasouj University |
| MNCN19119 | *Lanius senator senator* | F | Spain | Madrid, Villavieja de Lozoya | MNCN |
| MNCN19513 | *Lanius senator senator* | F | Spain | Madrid, Manjirón | MNCN |
| MNCN19601 | *Lanius senator senator* | F | Spain | Madrid, Puentes Viejas | MNCN |
| MNCN41114 | *Lanius senator senator* | F | Spain | Madrid, Horcajo de la Sierra | MNCN |
| woodchat 3 | *Lanius senator niloticus* | M | Iran | Kohgiluyeh and Boyer-Ahmad Province, Yasuj | Yasouj University |
| woodchat 4 | *Lanius senator niloticus* | M | Iran | Kohgiluyeh and Boyer-Ahmad Province, Yasuj | Yasouj University |
| woodchat 5 | *Lanius senator niloticus* | M | Iran | Kohgiluyeh and Boyer-Ahmad Province, Yasuj | Yasouj University |
| woodchat 6 | *Lanius senator niloticus* | M | Iran | Kohgiluyeh and Boyer-Ahmad Province, Yasuj | Yasouj University |
| IT2085 | *Lanius senator badius* | M | Italy | Sardinia, Cardedu | W.I.N.E.S. |
| IT2086 | *Lanius senator badius* | F | Italy | Sardinia, Cardedu | W.I.N.E.S. |
| IT2087 | *Lanius senator badius* | F | Italy | Sardinia, Ilbono-Elini | W.I.N.E.S. |
| NHMO-BI-15168/2 | *Lanius senator niloticus* | M | Israel | Eilat | NHMO |
| NHMO-BI-15169/2 | *Lanius senator niloticus* | F | Israel | Eilat | NHMO |
| NHMO-BI-15277/1 | *Lanius senator niloticus* | M | Israel | Eilat | NHMO |
| NHMO-BI-15274/2 | *Lanius senator niloticus* | F | Israel | Eilat | NHMO |
| MZB 2003-1092 | *Lanius senator senator* | M | Spain | Cervià de les Garrigues, Lleida - Les Garrigues | MZB |
| MZB 2011-0953 | *Lanius senator senator* | F | Spain | Catalunya | MZB |
| MZB 2011-1010 | *Lanius senator senator* | M | Spain | \ | MZB |
| MZB 2005-0946 | *Lanius senator senator* | M | Spain | Balaguer, Lleida - La Noguera | MZB |
| MCCI 3114 | *Lanius senator badius* | F | Italy | Sicily, Marettimo - Isola | MCC |
| MCCI 4629 | *Lanius senator senator* | M | Italy | Liguria, Savona | MCC |
| MCCI 4107 | *Lanius senator niloticus* | M | Armenia | Meghri | MCC |
| MCCI 4113 | *Lanius senator niloticus* | F | Armenia | Meghri | MCC |
| MCCI 4227 | *Lanius senator niloticus* | M | Armenia | Meghri | MCC |
| AY599852 | *Lanius senator senator* | \ | Italy | — | GenBank |
| AY599853 | *Lanius senator senator* | \ | Italy | — | GenBank |
| AY599854 | *Lanius senator niloticus* | \ | Israel | — | GenBank |
| AY599855 | *Lanius senator badius* | \ | Italy | — | GenBank |
| IPMB 10541* | *Lanius senator senator* | \ | France | southern France | GenBank |
| IPMB 12701* | *Lanius senator niloticus* | \ | Israel | — | GenBank |
| IPMB 8644* | *Lanius senator senator* | \ | Greece | Crete | GenBank |
| HQ996787 | *Lanius senator niloticus* | \ | Israel | — | GenBank |
| JF498788 | *Lanius senator niloticus* | \ | Iraq | Muhafazat Salah ad Din, Joint Base Balad | GenBank |
| JQ175216 | *Lanius senator senator* |  | Greece | Lesvos | GenBank |
| JQ175217 | *Lanius senator senator* | \ | Greece | Lesvos | GenBank |
| JQ175218 | *Lanius senator senator* |  | Greece | Lesvos | GenBank |

\*Voucher specimens

In order to achieve a broader assessment of geographic structure, we added to the molecular database Woodchat Shrike sequences from Greece, Iraq, France and Israel available on the GenBank and BOLD systems (i.e., COI: 4 sequences; *cytb*: 3 sequences; CR: 4 sequences; MYO: 4 sequences; Table 1 and S2, Figure 1). When missing, the putative subspecies of GenBank sequences and museum specimens were inferred based on the sampling site, collection date and the currently recognized geographical distribution of the taxa. The final dataset includes representative individuals from localities encompassing the breeding distribution of the three currently recognized subspecies (i.e., *L. s. senator, L. s. badius*, and *L. s. niloticus*). We selected four mitochondrial markers to investigate the genetic structure of the species: NADH dehydrogenase subunit 2 (ND2), cytochrome b (*cytb*), cytochrome c oxidase I (COI), and a fragment of the control region (CR). Additionally, for a subset of specimens we sequenced the nuclear intron 2 of myoglobin (MYO), and introns 6– 7 of the ornithine decarboxylase (ODC) genes (Table S2).

### DNA extraction, Amplification and Sequencing

Total genomic DNA was isolated with the NucleoSpin Tissue kit (Macherey-Nagel, Düren, Germany), following the manufacturer’s protocols. Afterward, we amplified and sequenced the selected loci, using a combination of published primers following molecular procedures as described in Pellegrino et al. (2017). Primer details and annealing temperatures are reported in Table S3. When the first round of amplification or sequencing failed, and for the amplification of nuclear loci, we performed PCRs by using the with puReTaq Ready-To-Go PCR beads (Amersham Bioscience, Freiburg, Germany) in a 25 µL reaction according to the manufacturer’s given instructions.

After verifying amplicon occurrence and length through agarose gel electrophoresis, the PCR products were cleaned with ExoSAP-ITA PCR Product Cleanup Reagent (Thermo Fisher Scientific, Waltham, Massachusetts, USA). The sequencing reaction was performed with an ABI 3730xl DNA Analyzer by Macrogen Europe Inc. (Amsterdam, The Netherlands) using the same PCR primer pairs, except for ND2 H1064 which was replaced by the internal forward primer L347.

### Haplotype analysis

Electropherograms were visualized and modified with Bioedit 7.2 (Hall 1999), and for nuclear markers, the heterozygous sites were corrected using the nucleotide IUPAC code. Forward and reverse runs were aligned with the ClustalW algorithm (Larkin et al. 2007) with default settings.

All the nucleotide sequences generated in this study were submitted to GenBank (accession numbers OD991799 - OD992005, Table S2).

Aligned nucleotide sequences were collapsed into unique haplotypes using FaBox 1.5 (Villesen 2007) for each single marker and for the concatenated dataset. For each sample, the nucleotide sequences of the four analyzed mitochondrial markers were concatenated to create a unique dataset.

In order to investigate the frequency and geographic distribution of mtDNA haplotypes, Median Joining networks, encompassing all the investigated *Lanius senator* populations (Figure 1), were built using the median-joining algorithm implemented in PopART 1.7 (Leigh and Bryant 2015). Haplotype networks were generated for each single mtDNA marker (Figure S1) and for the concatenated multilocus dataset (Figure 2).

**Figure 2.**
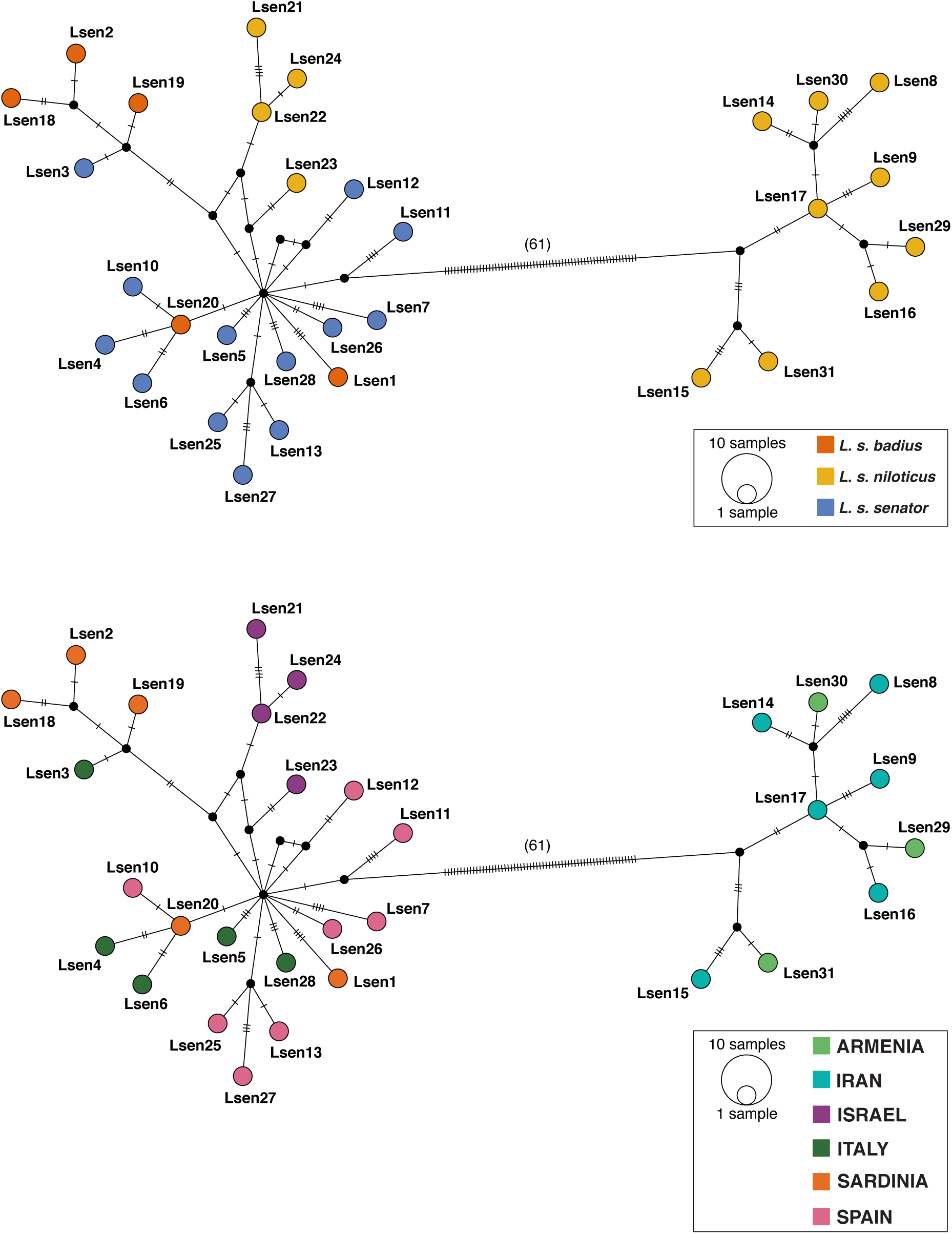
Haplotype median-joining networks of the concatenated mtDNA dataset. Figure 2a shows haplotypes network based on putative subspecies designations (see inset), while Figure 2b is based on sampling locality (see inset).

### Phylogenetic analyses and molecular dating

Mitochondrial gene tree reconstruction was performed using Bayesian Inference (BI) implemented in MrBayes 3.2 (Ronquist et al. 2012), for haplotypes belonging to both the single markers and the concatenated dataset. Available sequences were retrieved for *L. senator* and other *Lanius* species from GenBank NCBI and were added to the original dataset in order to increase the information and to root the trees (Table S2, Figure 3). The best-fitting nucleotide substitution model was inferred using JModelTest 2.1 (Posada 2008), under the corrected Akaike Information Criterion (AIC), and used as follow: GTR+G for ND2, HKY+I for *cytb*, GTR+I+G for COI, GTR+G for CR, and a partition scheme, corresponding to the length of the sequences for each locus, was set to analyze the concatenated data, applying the previously determined models. Two independent runs were performed by setting 4 Metropolis-coupled MCMC, 3 heated and 1 cold, with temperature command equal to 0.2. A total number of 2^*^10^6^ iterations were performed, with sampling every 1000 iterations; 25% of the samples were discarded as burn-in. We ensured that the standard deviation of split frequencies converged towards zero and that the potential scale reduction factor (PSRF) was reasonably close to 1.0 for all the parameters. Furthermore, we checked the outcome files in Tracer 1.7 (Rambaut et al. 2018) to make sure the posterior distribution of the runs converged and that the ESS was higher than 200 for meaningful parameter estimations.

**Figure 3.**
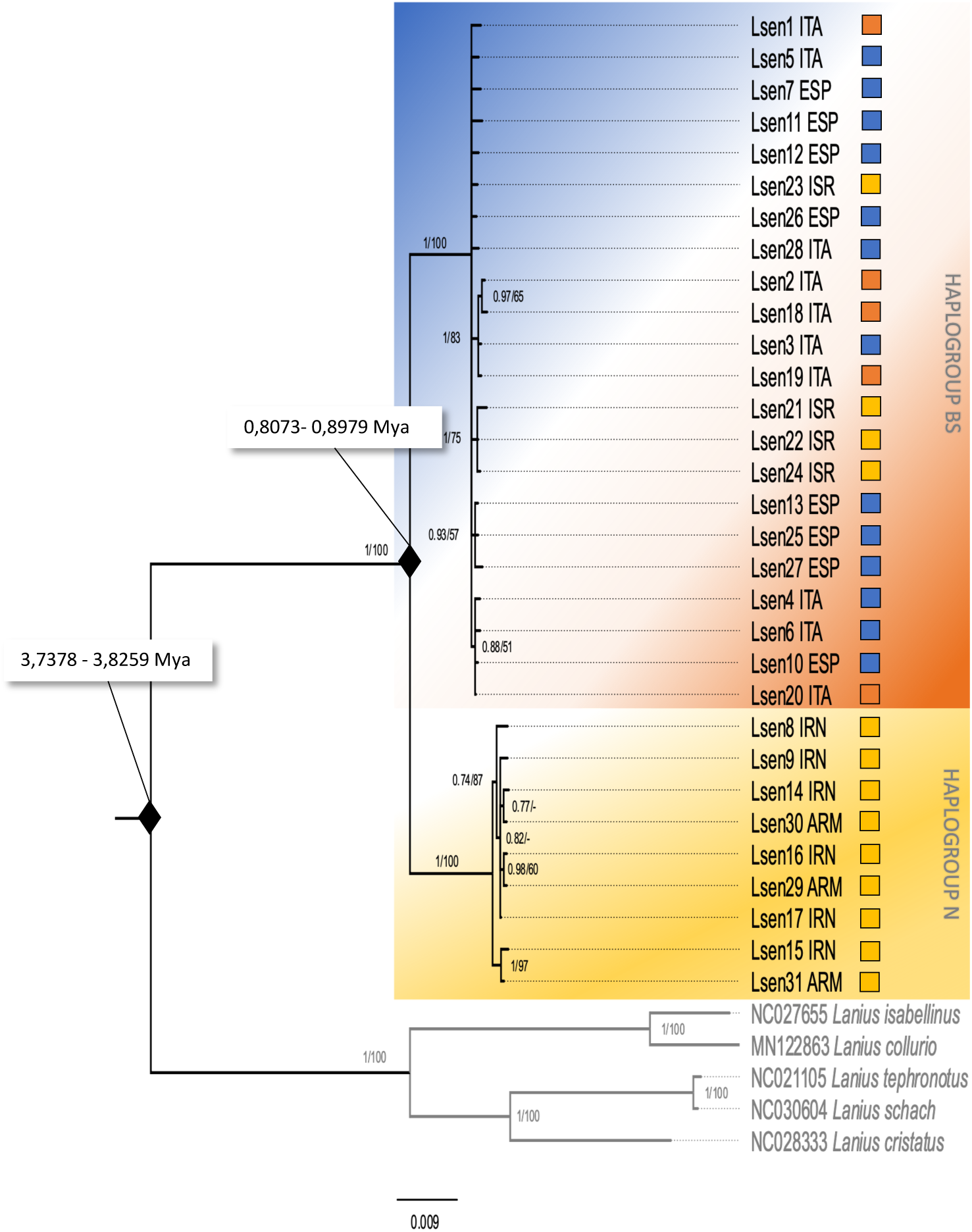
Bayesian Inference and Neighbor-Joining phylogenetic tree of the concatenated mtDNA haplotypes. Numbers on the nodes refer to BI posterior probabilities (left) and to bootstrap support values of the NJ analysis (right) using the p-distance substitution model. Thresholds for both BI posterior probabilities and bootstrap support values were set at ≥ 50%; dashes represent values below that threshold. The colours shown here mimic those used in Figure 1, such that the orange-blue background corresponds to the *badius/senator* (BS) haplogroup, while the yellow background represents the *niloticus* (N) haplogroup. Squares to the right of each individual define the putative subspecies attributed to the individual, based on geography and morphology.

In addition to the BI reconstruction, we also performed a Neighbor-Joining (NJ) (Saitou and Nei 1987) analysis for the mtDNA concatenated dataset employing Mega X (Kumar et al. 2018), and using p-distance. The topology of the trees was visualized with FigTree 1.4 (Rambaut 2008). Sequences from different species of *Lanius* genus were added as outgroups in the five gene? dataset, GenBank ID and species details are indicated in figures 3, S2a-d. In addition, one *L. senator* available sequence on GenBank from Gambia was aligned in the dataset of ND2 sequences in order to investigate its position in a phylogenetic tree.

For the nuDNA markers analysis, we used PHASE 2.1 (Stephens et al. 2001) as implemented in DNASP 5.10 (Librado and Rozas 2009) to infer heterozygous sites and the corresponding alleles. Two runs were performed, one with default settings and the second with 1000 iterations and 100 of burn-in. We used a posterior probability threshold of 0.9 to determine the most probable haplotype for each nuclear sequence and we removed the individuals that did not satisfy this threshold; no significant differences were found between the two runs. As an alternative to concatenation, we performed a Bayesian concordance analysis (BCA) for nuclear haplotypes tree, employing BUCKy 1.4 (Larget et al. 2010). We first created two different gene trees, one for MYO and one for ODC intron sequences, running a single run in MrBayes and setting 4 chains, 1,000,000 iterations, sampled every 1000 generations. The best nucleotide substitution models inferred by JModelTest were HKY+I for both the nuclear markers. Subsequently, we summarized the generated trees, and the first 25% of the trees were discarded as burn-in. The resulting output was processed in BUCKy to create a concordance tree and to calculate the concordance factors (CF) for clades. We used four different settings, each one of them was performed with four independent runs, four Metropolis-coupled MCMC (3 heated and one cold), but different a priori level of discordance among loci (α), that was set at 0.1, 1, 10, and 100 respectively; no significant dissimilarities were found in CFs between the settings, so we report CFs for the α value of 100 (Figure S4). Tree topology was visualized in FigTree 1.4.

Divergence time between the main clades was estimated through a Bayesian approach in Beast 1.10 (Suchard et al. 2018) using ND2 and *cytb* markers. We incorporated best fit models for each gene and applied the lineage substitution rate of 0.014 per lineage/million years for *cytb* and 0.029 per lineage/million years for ND2 with standard deviation of 0.001 and 0.0025 respectively (Lerner et al. 2011). The analyses were performed using a strict molecular clock with fixed mutation rates and the tree prior constant Yule’s speciation. A Yule process speciation prior was implemented in each analysis. Three separate MCMC analyses were run for 100,000,000 generations with parameters sampled every 10,000 steps, with a 10% burn-in, producing 10,000 trees each. The independent runs were combined using LogCombiner v.1.6.1 (Drummond et al. 2012). Tracer v.1.7 was used to analyse the convergence of the posterior distribution in the three runs and to measure the effective sample size of each parameter (all >200). Tree topologies were assessed using TreeAnnotator v.2.6.3 and FigTree v.1.4.

For each haplogroup, standard genetic indices, such as haplotype diversity, polymorphic sites, and nucleotide diversity, were calculated with DNASP 5.10. We also estimated neutrality parameters Fu’s Fst and Tajima’s D test with Arlequin 3.5 (Excoffier and Lischer 2010). We performed 1000 simulated samples with a threshold of 0.9 to obtain these values.

### Genetic distances

Orthologous ND2, COI, *cytb* and CR sequences, belonging to 13 *Lanius* species, were downloaded from GenBank-NCBI. After alignment (conducted with MAFFT 7.110; (Katoh and Standley 2013); using the E-INS-I option), sequences showing insertions/deletions (with the only exception of CR), that were missing > 1% of sites or that were overlapping in length less than 75% with the region amplified and sequenced for the Italian samples, were discarded. Overall, a total of 35 ND2, 290 COI, 309 *cytb*, and 29 CR sequences were retrieved from GenBank NCBI and combined with the 34 sequenced *Lanius senator* samples to constitute a unique dataset for each mtDNA marker (see Table S4).

We calculated the uncorrected pairwise genetic distances (p-distance) between and within groups (i.e., the species and the two haplogroups of *L. senator*) using MEGA X (Kumar et al. 2018). A condensed list of these values (i.e., average and range) per marker and grouping (i.e., *L. senator*, the two haplogroups of *L. senator*, and all the species belonging to the genus *Lanius* except for *L. senator*) are reported in Table 2, while the average p-distance values for each species are reported in Table S5.

**Table 2.**
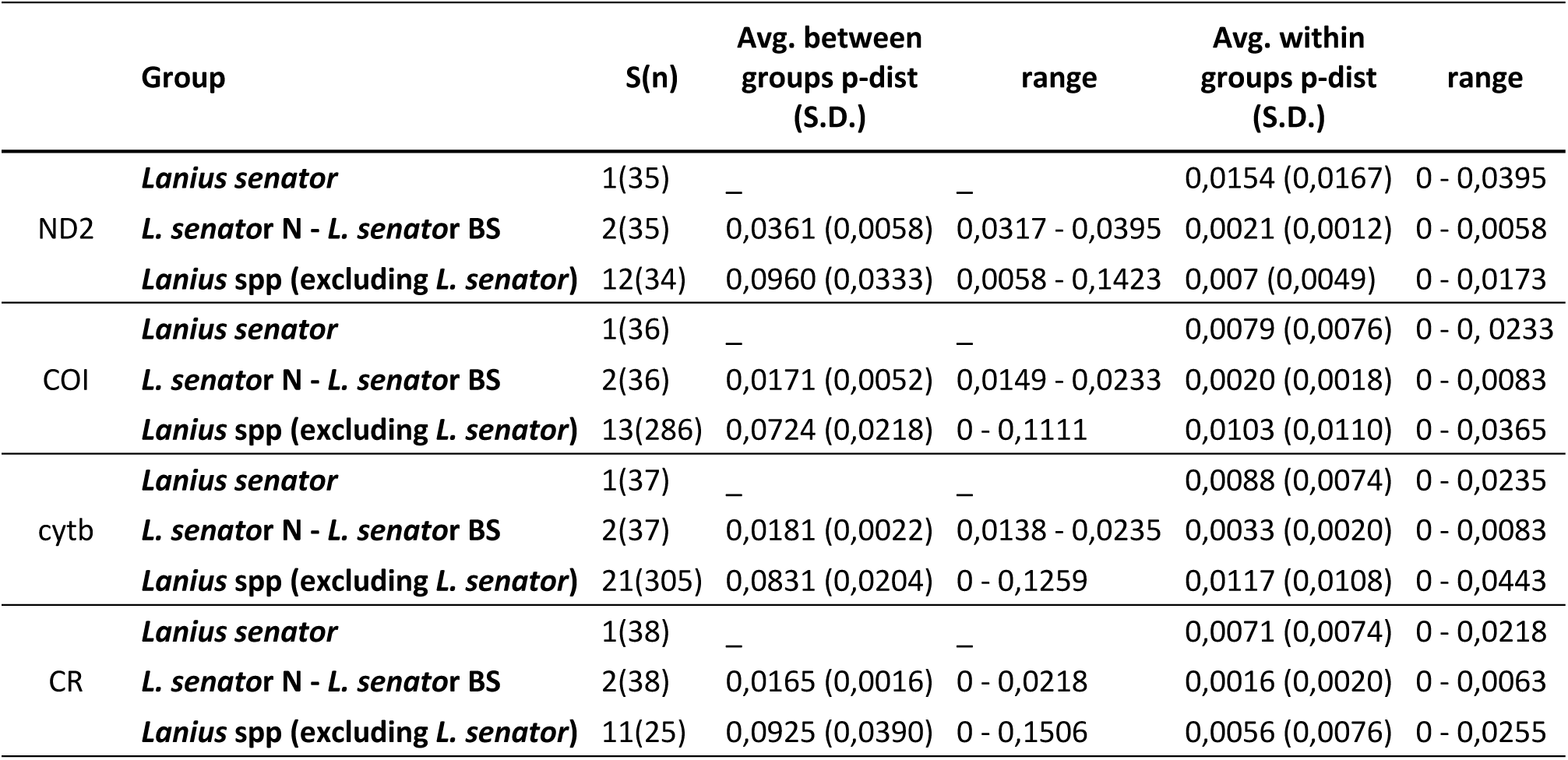
Genetic diversity indices calculated on mtDNA loci, concatenated mtDNA dataset, and nuDNA loci. ^*^ (p < 0.05), ^**^ (p < 0.01).

## Results

DNA was successfully extracted from all samples, and we obtained 34 sequences of ND2, *cytb*, and CR and 32 of COI for a total of 997 bp of ND2, 660 bp of COI gene, *cytb* 965 bp and 327 bp of CR. mtDNA sequences were concatenated to obtain 2924 bp sequences. Moreover, we sequenced MYO (693 bp) and ODC (489 bp) nuclear genes in a subsample of 14 and 16 individuals, respectively (Table S2).

All identified mtDNA haplotypes were new, except for one haplotype of COI (lsenCOI_3 corresponding to the GenBank record JQ175216 and JQ175218 sampled in Greece) and one of CR (Lsencr_1 with AY599853 and AY599855 from Italy recorded as *L. s. senator* and *L. s. badius* respectively).

After phasing the sequences of heterozygous individuals, we identified a total of eight haplotypes in MYO with eight variable sites and 10 haplotypes with nine variable sites in ODC sequences. The recognized haplotypes were new except for two haplotypes of MYO: LsenMyo*_*1, the most frequent haplotype identified in individuals from Spain, Italy and Israel was recovered also in three individuals from GenBank sampled in France, Israel and Greece; and LsenMyo_2 identified from Sardinia and Armenia which corresponds to a GenBank record from Israel (Table S2).

### Haplotype analyses

The haplotypes median joining network (Figures 2a and 2b) based on the concatenated mtDNA dataset highlighted two different clusters separated by 61 mutations. The first cluster, containing the largest number of haplotypes and samples, includes *L. s. senator* individuals from Italy and Spain, *L. s. badius* individuals from Sardinia and presumed *L. s. niloticus* from Israel. The second cluster includes all the individuals from Armenia and Iran, which are all ascribed to *L. s. niloticus*.

Haplotype networks calculated on single markers (Figure S1) confirmed the two main clusters, with the same structure. All the haplotypes within each haplogroup differed by only one to three mutations. These single marker haplotype networks showed that individuals identified as different subspecies can share haplotypes, for instance LsenND2_3 was shared by both *L. s. badius* and *senator*, and LsenCOI_3 included individuals attributed to all three subspecies. The most frequent haplotype, identified for ND2, was LsenND2_4 shared by six individuals from Italy, Israel and Spain and ascribed to *L. s. niloticus* or *L. s. senator*, while LsenCytb_6 was shared by three individuals from Italy that were morphologically identified as *L. s. senator* or *L. s. badius* (Figure S1).

### Phylogenetic analyses and molecular dating

Neighbour-joining and Bayesian inference trees constructed using the concatenated mtDNA dataset, as well as all individual loci datasets each recovered two well supported (Bayesian posterior probability = 1.0) clades (Figure 3, Figure S2 a-d), which corresponded with the haplotype network results. One monophyletic clade (N) included individuals ascribed to the subspecies *L. s. niloticus*, from Iran and Armenia. The second clade (BS) included all individuals from Israel, Greece, Italy, Sardinia, Sicily, France and Spain, and thus encompasses birds classified as belonging to the three recognized subspecies of Woodchat Shrike (*L. s. senator, L. s. badius L. s. niloticus)*. A GenBank sequence of a Woodchat Shrike from Gambia, part of the wintering range of the subspecies *L. s. senator*, clustered within the BS haplogroup. Bayesian inference trees obtained for each individual gene (Figure S2a-d) highlighted that some haplotypes were restricted to unique geographical populations, and in particular the *cytb* phylogeny identified a moderately well-supported clade (0.85 estimated bayesian posterior probability), comprising all samples from Israel (Figure S2c).

The phylogenetic trees based on nuclear markers revealed poorly structured clades (Figure S3 a-b), and no clear geographical differentiation. Individuals from Armenia and Iran (identified as *L. s. niloticus*, clade N in the mtDNA analyses) showed the same haplotype or clustered with individuals ascribed to the other two subspecies. The Bayesian concordance analysis (BCA) showed a clade with a low concordance including only individuals from Armenia and Iran (CF = 0,018) and a clade with a relatively high concordance (CF=0,5) grouping exclusively individuals from all other sampling areas.

Our BEAST analysis on the more variable markers *cytb* and ND2 yielded two similar time-calibrated trees for *Lanius senator*. The divergence time was estimated for the *Lanius senator* clade at 3.7378 million years ago (Mya) (height 95% HPD: 2.8344-4,8286) and 3.8259 Mya (height 95% HPD:2.7422 - 4.7716) for ND2 and *cytb* respectively. The divergence time from BS and N haplogroups was estimated at 0.8073 Mya (height 95% HPD: 0.5496 - 1.1279) by ND2 and 0,8979 (height 95% HPD: 0.5761 - 1.2595) by *cytb* (Figure 3).

In order to evaluate the genetic distances between and within *Lanius* species, we constructed four datasets, one for each mtDNA locus, encompassing all the available sequences from GenBank, BOLD systems and from this study. The resulting uncorrected pairwise genetic distances (p-distance) are shown in Table 2. The average p-distances between *Lanius* species (excluded *L. senator*) ranged from 0.0724 (± 0.0218) in COI to 0.0960 (± 0.0333) in *cytb*. Average distance between the *Lanius senator* haplogroups N and BS ranged from 0.0165 (± 0.0016) in CR to 0.0361 (± 0.0058) in ND2. Average distances within *Lanius* species (excluded *L. senator*) ranged from 0.007 (± 0.0049) in ND2 and 0.0117 (± 0.0108) in *cytb*. Within *Lanius senator*, the p-distances showed the highest value for ND2 (0.0154 ± 0.0167), and both ND2 and CR had higher values than the p-distances within the entire *Lanius* genus. Average p-distances estimated within the N and the BS groups were highest for *cytb* (0.0033 SD: 0.0020) and lowest for CR (0.0016 ± 0.0020).

Estimated genetic diversity indices calculated in each clade and in the total dataset for each analyzed marker are shown in Table 3. In mtDNA genes haplotype diversity (*h*) ranged from 0.962 (± 0.016) for *cytb* to 0.661 (± 0.075) for CR region. The highest value of *h* was found in the haplogroup N for ND2 (0.972 ± 0.064), and the lowest value was for CR (0.222 ± 0.166), also in haplogroup N (Table 3). A significantly negative Tajima’s D value was observed for the BS haplogroups at all mtDNA markers, and a significantly negative Fu & Li’s D was observed at each of the four mtDNA markers across all haplogroups, with the exception of CR and cytb markers for the N haplogroup (Table 3). Haplotype diversity calculated on MYO and ODC sequences were 0.831 (± 0,045) and 0,613 (± 0,088) respectively; Tajima’s D and Fu & Li’s D test showed negative and significative values in the MYO dataset.

**Table 3.**
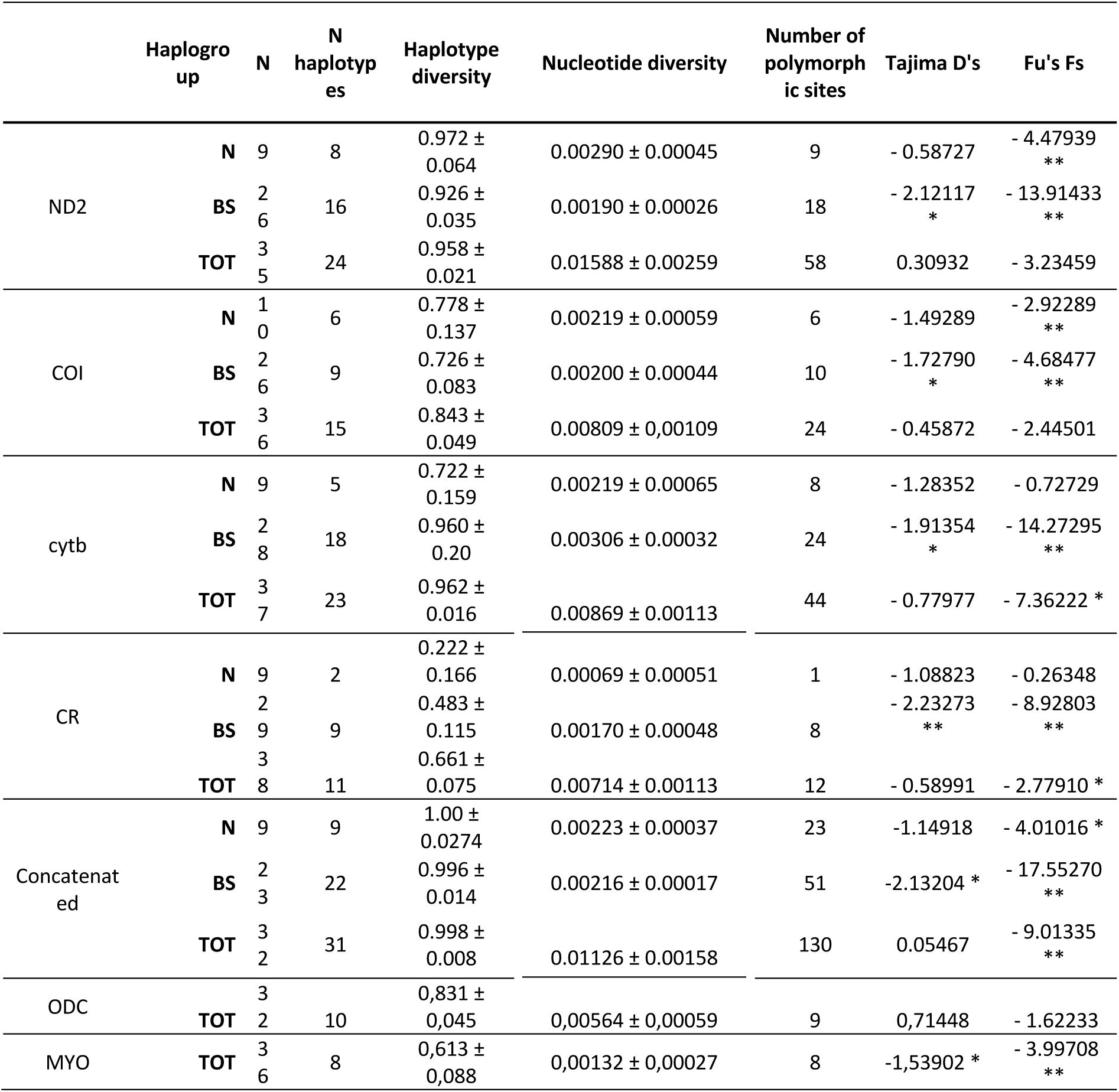
Genetic distances (p-distance) dataset between and within different groups: within *Lanius senator*, between and within *L. senator* haplogroups N and *L. senator* haplogroups BS; *Lanius* spp. (excluding *L. senator*). Distances were estimated for all the mtDNA loci dataset (Table S4). Column S(n) includes the number of considered group/species and, within brackets, the overall number of considered sequences.

## Discussion

Our investigation of phylogeographic variation in the Woodchat Shrike showed unexpected results when considering the current state of knowledge regarding the three recognized subspecies’ distributional ranges and plumage diagnostic features.

Mitochondrial tree reconstruction and haplotype networks showed that our samples were not divided into three clades/haplogroups, as we initially hypothesized based on the three sampled subspecies but were instead assigned to two strongly supported clades. One clade (N) contained individuals from Iran and Armenia, which correspond to the eastern portion of the putative *L. s. niloticus* breeding range (Figure 1). The second clade (BS) grouped all the individuals from Italy, Spain, France, Greece, as well as those from Israel and Sardinia. Individuals from Israel were originally assigned as *L. s. niloticus* on the basis of plumage characteristics, sampling sites (see Figure 1) and dates (see Shirihai 1996), while individuals from Sardinia were referred to *L. s. badius* based on plumage and sampling location (see Figure 1). All other individuals in this clade were identified as *L. s. senator*. Genetic distance between the two haplogroups ranged from 1.7 and 3.6 % depending on the molecular locus; these values are in the range of interspecific and intraspecific avian variability (Barrowclough et al. 2016).

Conversely to what we found for mitochondrial markers, the nuclear markers MYO and ODC showed low variability and did not reveal any clear geographical structure. Such discordance could be due to incomplete lineage sorting, ghost speciation or asymmetrical sexual dispersion (Toews and Brelsford 2012). In this context, the mito-nuclear discordance we found here, matches with findings in other Mediterranean populations of Passeriformes such as *Muscicapa tyrrhenica* or *Certhia familiaris* (Pons et al. 2016, 2019), and in general this discordance is prevalent in different animals’ system (Toews and Brelsford 2012). A genome-wide sequencing approach could be useful to highlight genetic variation within such species to clarify the processes underlying the lack of structuring in nuclear genes (Calderón et al. 2016; Ottenburghs et al. 2019).

A number of phylogenetic studies on *Lanius* species (Gonzalez et al. 2008; Klassert et al. 2008; Fuchs et al. 2011; Olsson et al. 2013; Pârâu et al. 2019) have revealed that genetic data are often inconsistent with morphological characteristics (Gill et al. 2021), as well as already found in other avian species (e.g., *Motacilla* spp. Li et al. 2016; Drovetski et al. 2018b; *Emberiza striolata* Schweizer et al. 2017).

Phylogeographic studies have identified multiple clades in three shrike species, *L. excubitor* and *L. collurio*, although these did not show the same geographical distribution of *L. senator* as the clades in the other species, nor is there an indication of clade diversification resulting from possible isolation in glacial refugia in either species (Olsson et al. 2010; Pârâu et al. 2019). In the case of *L. senator*, we recovered two haplogroups within the breeding range of the species showing a widely distributed western haplogroup and a narrowly distributed eastern one. These distributions could support diversification of the two clades in two different areas, the Southern Caucasus in the case of *L. s. niloticus* and Iberia, Greece and Italy for *Lanius s. senator and L. s. badius*. Future studies could highlight whether the Maghreb region in North Africa could have played a role as a glacial refugium for this species. The estimated origin of ∼0.85 Mya for these two clades occurred during the Mid-Pleistocene Climate Transition (1.2 - 0.7 Mya), a shift in paleoclimatic periodicity from 41- to 100-kyr cycles, which led to major changes in global ice volume, sea level, and ocean temperature (Ruddiman et al. 1989), as well as an increase in aridity in Africa and the Arabian Peninsula (deMenocal 2004). The subsequent habitat reductions and changes could have divided and separated populations both in the wintering and in breeding areas, thereby resulting in genetic diversification. A similar divergence time between clades and a western versus eastern Palearctic distribution has been found in different bird species adapted to arid climates such as *Galerida* spp. (Guillaumet et al. 2008) and *Chlamydotis* spp. (Korrida and Schweizer 2013). Fuchs et al. (2019) calculated the probability of the ancestral area for each *Lanius* species. Given their results, Africa (75%) and Europe (15%) were identified as ancestral areas for *L. senator*, which clustered in a clade of African origin. Therefore, the current distribution of *L. senator* is probably a consequence of an expansion from Africa, which may have begun with the colonization of the Arabian Peninsula. This colonization could have been facilitated by the presence of the Afro-Arabian land bridge in the Mid-Pleistocene (1 mya, Bosworth et al. 2005). This connection has been implicated in colonization patterns for different terrestrial vertebrate lineages (Pook et al. 2009; Portik and Papenfuss 2015) including birds (Voelker et al. 2016). Considering the distribution of haplogroup N in the breeding range of *L. senator*, the Caucasian glacial refugium might have played an important role in preserving the observed genetic differences, as already shown for several avian species (Perktaş et al. 2015; Drovetski et al. 2018a; Pavia et al. 2021).

Surprisingly, in the western clade, our mtDNA and nuDNA data did not allow us to genetically distinguish between the two putative subspecies, *L. s. senator* and *L. s. badius*. These taxa display both an allopatric distribution and distinctive characteristics, to include bill size and shape, wing formula and plumage features; moreover, they overwinter in putatively different ranges (Small and Walbridge 2005). The morphological differentiation between the two taxa could have occurred very recently and as such are not yet distinguishable through molecular markers (or at least those we analysed). The differentiation could be due to a refugial isolation during the Last Glacial Maximum or evolutionary diversification mediated by insular isolation. Differently from *Lanius senator*, well-defined Mediterranean lineages were found in different migratory birds to include *Muscicapa tyrrhenica, Curruca sarda* and the *Curruca cantillans* complex (Pons et al. 2016; Zuccon et al. 2020; Nespoli et al. 2021), as well as several mammals, butterflies, and reptiles (Grill et al. 2007).

The incoherence between genetics and morphology could be due to an early stage of divergence with gene flow, or different selective conditions in the species range or sexual selection. Morphological and behavioral traits could play a decisive role in taxa differentiations (Nwankwo et al., 2017) and for this reason they should be investigated and considered along with molecular markers. Moreover, our results showed also that this lack of genetic distinction also concerns the putative subspecies *L. s. rutilans*, which some authors recognize as valid (Table S1). However, all the Iberian individuals, including those from the described range for *rutilans* are part of the BS haplogroup, where there is no Iberian-distributed substructure.

Negative and significant values of Tajima’s D and Fu & Li’s D test in our results indicate a population expansion, these results could be affected by the wide distribution of the utilized samples set that comprises the much of the breeding range or these values could be indicative of a past population expansion, which is common in migrating species during post glacial expansion (Fahey et al. 2012). The current status of the species (Near threatened, BirdLife International 2021) makes it even more urgent to deepen our general knowledge of the genetic characteristics of the different populations. In fact, declining local populations could increase the risk of genetic consequence due to their small size (Pertoldi et al. 2021).

Lastly, we want to draw attention to the morphological identification of *L. s. niloticus*. This subspecies is often reported on minor (very small) Italian islands during the migration season (Brichetti and Fracasso 2007). Moreover, some of the individuals from Israel used in this study, identified as *niloticus*, clustered with *badius/senator*. The weak distinctive morphological characteristics could be due to the absence of definition of unique characteristics or hybridization between the two subspecies. Future work will be focused on determining whether the two subspecies hybridize in contact zones. *The subspecies breeding distributions should be investigated thoroughly, particularly in the taxonomically uncertain ranges in Turkey (Roselaar 1995)*, where Woodchat Shrike morphology is characterized by a gradient from west to east *(i*.*e*., *eastern L. s. senator* tend to resemble *L. s. niloticus*; Shirihai and Svensson 2018).

On the whole, our study showed that morphological characteristics alone can no longer be considered decisive for identifying *Lanius senator* subspecies with certainty and lays the groundwork for the development of a new diagnostic set of characters.

Although the present work has brought new insights into the phylogeographical aspects of *Lanius senator*, a second step will have to focus on sampling wintering areas and adding individuals from the Maghreb and Balearic Islands in order to have a more complete framework.

A final consideration concerns the conservation implications of Woodchat Shrike in a context of populations declining in different parts of its range (e.g., Italy, Brichetti and Grattini, 2017) and the recent worsening of its conservation status, having been shifted from Least Concern to Near Threatened (BirdLife International 2021). The phylogeographic data highlighted two highly distinct and unique clades that both deserve particular attention in conservation efforts. New genetic data that include samples from the Balearic Islands and across the wintering range could help to understand whether *L. s. badius* can be considered a valid subspecies or an evolutionarily significant unit (ESU) characterized by distinctive behavioural and phenotypic traits that may deserve conservation attention.

## Acknowledgements

We are grateful to the curators and directors of the following Museums and Collections for providing samples: Isabel Rey and Beatriz Alvarez Dorda (MNCN, Madrid, Spain); Lars Erik Johannessen (NHMO, Oslo, Norway); Javier Quesada (MZB, Barcelona, Spain). We wish to thank ISPRA (dr. Lorenzo Serra), Regione Sardegna, Calabria and Abruzzo to give the permits to collect the biological samples. We also thank Andrea Romano for providing helpful comments. Finally, we thank for the field support Gianluca Congi, Gloria Ramello and Elisa Mancuso. This is publication number XXXX of the Biodiversity Research and Teaching Collections at Texas A&M University and publication number 10 of the W.I.N.E.S. collaborative research group.

## Additional Information

### Data Accessibility

All specimen data are accessible on GenBank (Accession numbers OD991865-OD992005).

## Supplementary material

### Tables

- Supplementary material S1. Table of described subspecies of *Lanius senator*
- Supplementary material S2. Table of samples used in this study with specific information on sampling locality and date, type of specimen, sex/age, collocation of sample material and collectors’ contacts. We also report the haplotype attributed to the sample, the resulting haplogroup and GenBank accession number.
- Supplementary material S3. PCR amplification details, primer used and thermocycling conditions for each locus investigated.
- Supplementary material S4. List of GenBank sequences used to estimate the genetic distances reported in test and in table S5.
- Supplementary material S5. Genetic distances for each mtDNA dataset, calculated between and within every *Lanius spp*., including our *badius/senator* haplogroup (BS) and *niloticus* haplogroup (N).

### Figures

- Figure S1: Haplotypes’ median-joining networks of each mtDNA dataset (*i*.*e*., ND2, COI, *cytb*, CR), based on putative subspecies.
- Figure S2a: Bayesian Inference phylogenetic tree of the ND2 mtDNA haplotypes. Numbers on the nodes refer to BI posterior probabilities, with threshold set at ≥ 50%. The colours shown here recall the ones used in Figure 1 and 3.
- Figure S2b: Bayesian Inference phylogenetic tree of the COI mtDNA haplotypes. Numbers on the nodes refer to BI posterior probabilities, with threshold set at ≥ 50%. The colours shown here recall the ones used in Figure 1 and 3.
- Figure S2c: Bayesian Inference phylogenetic tree of the *cytb* mtDNA haplotypes. Numbers on the nodes refer to BI posterior probabilities, with threshold set at ≥ 50%. The colours shown here recall the ones used in Figure 1 and 3.
- Figure S2d: Bayesian Inference phylogenetic tree of the CR mtDNA haplotypes. Numbers on the nodes refer to BI posterior probabilities, with threshold set at ≥ 50%. The colours shown here recall the ones used in Figure 1 and 3.
- Figure S3a: Bayesian Inference phylogenetic tree of the ODC nuDNA haplotypes. Numbers on the nodes refer to BI posterior probabilities, with threshold set at ≥ 50%. The colours shown here recall the ones used in Figure 1 and 3.
- Figure S3b: Bayesian Inference phylogenetic tree of the MYO nuDNA haplotypes. Numbers on the nodes refer to BI posterior probabilities, with threshold set at ≥ 50%. The colours shown here recall the ones used in Figure 1 and 3.
- Figure S4: Bayesian Concordance Analysis tree of ODC and MYO dataset. Numbers on the nodes refer to CFs values, with threshold set at ≥ 50%. The colours shown here recall the ones used in Figure 1 and 3

