## Supplemental Material_Figures for "Phylogeography of *Lanius senator* reveals conflicts between alpha taxonomy, subspecies ranges and genetics"

**Captions:**

**Figures.**

- **Figure S1:** Haplotypes' median-joining networks of each mtDNA dataset (*i.e.,* ND2, COI, *cytb*, CR), based on putative subspecies.
- **Figure S2:** Bayesian Inference phylogenetic trees of mitochondrial haplotypes
  - Figure S2a: Bayesian Inference phylogenetic tree of the ND2 mtDNA haplotypes. Numbers on the nodes refer to BI posterior probabilities, with threshold set at ≥ 50%. The colours shown here recall the ones used in Figure 1 and 3.
  - Figure S2b: Bayesian Inference phylogenetic tree of the COI mtDNA haplotypes. Numbers on the nodes refer to BI posterior probabilities, with threshold set at ≥ 50%. The colours shown here recall the ones used in Figure 1 and 3.
  - Figure S2c: Bayesian Inference phylogenetic tree of the *cytb* mtDNA haplotypes. Numbers on the nodes refer to BI posterior probabilities, with threshold set at ≥ 50%. The colours shown here recall the ones used in Figure 1 and 3.
  - Figure S2d: Bayesian Inference phylogenetic tree of the CR mtDNA haplotypes. Numbers on the nodes refer to BI posterior probabilities, with threshold set at ≥ 50%. The colours shown here recall the ones used in Figure 1 and 3.
- **Figure S3:** Bayesian Inference phylogenetic tree of nuclear haplotypes
  - Figure S3a: Bayesian Inference phylogenetic tree of the ODC nuDNA haplotypes. Numbers on the nodes refer to BI posterior probabilities, with threshold set at ≥ 50%. The colours shown here recall the ones used in Figure 1 and 3.
  - Figure S3b: Bayesian Inference phylogenetic tree of the MYO nuDNA haplotypes. Numbers on the nodes refer to BI posterior probabilities, with threshold set at ≥ 50%. The colours shown here recall the ones used in Figure 1 and 3.
- **Figure S4**: Bayesian Concordance Analysis tree of ODC and MYO dataset. Numbers on the nodes refer to CFs values, with threshold set at ≥ 50%. The colours shown here recall the ones used in Figure 1 and 3 in the main text.

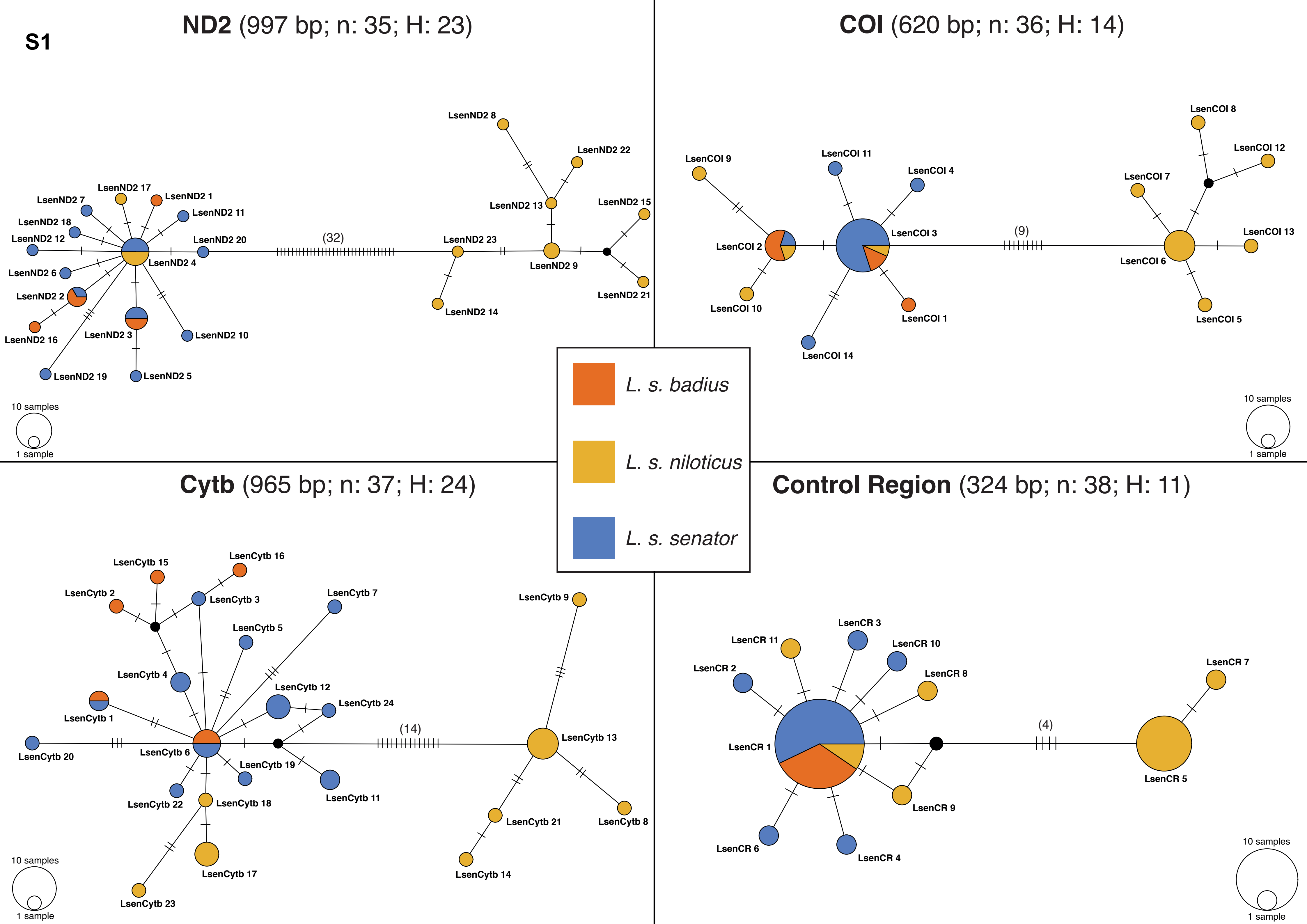
S1
ND2 (997 bp; n: 35; H: 23)
COI (620 bp; n: 36; H: 14)
Cytb (965 bp; n: 37; H: 24)
Control Region (324 bp; n: 38; H: 11)
L. s. badius
L. s. niloticus
L. s. senator
LsenND2 1
LsenND2 2
LsenND2 3
LsenND2 4
LsenND2 5
LsenND2 6
LsenND2 7
LsenND2 8
LsenND2 9
LsenND2 10
LsenND2 11
LsenND2 12
LsenND2 13
LsenND2 14
LsenND2 15
LsenND2 16
LsenND2 17
LsenND2 18
LsenND2 19
LsenND2 20
LsenND2 21
LsenND2 22
LsenND2 23
(32)
LsenCOI 1
LsenCOI 2
LsenCOI 3
LsenCOI 4
LsenCOI 5
LsenCOI 6
LsenCOI 7
LsenCOI 8
LsenCOI 9
LsenCOI 10
LsenCOI 11
LsenCOI 12
LsenCOI 13
LsenCOI 14
(9)
LsenCytb 1
LsenCytb 2
LsenCytb 3
LsenCytb 4
LsenCytb 5
LsenCytb 6
LsenCytb 7
LsenCytb 8
LsenCytb 9
LsenCytb 11
LsenCytb 12
LsenCytb 13
LsenCytb 14
LsenCytb 15
LsenCytb 16
LsenCytb 17
LsenCytb 18
LsenCytb 19
LsenCytb 20
LsenCytb 21
LsenCytb 22
LsenCytb 23
LsenCytb 24
(14)
LsenCR 1
LsenCR 2
LsenCR 3
LsenCR 4
LsenCR 5
LsenCR 6
LsenCR 7
LsenCR 8
LsenCR 9
LsenCR 10
LsenCR 11
(4)
10 samples
1 sample

**S2 (a)**

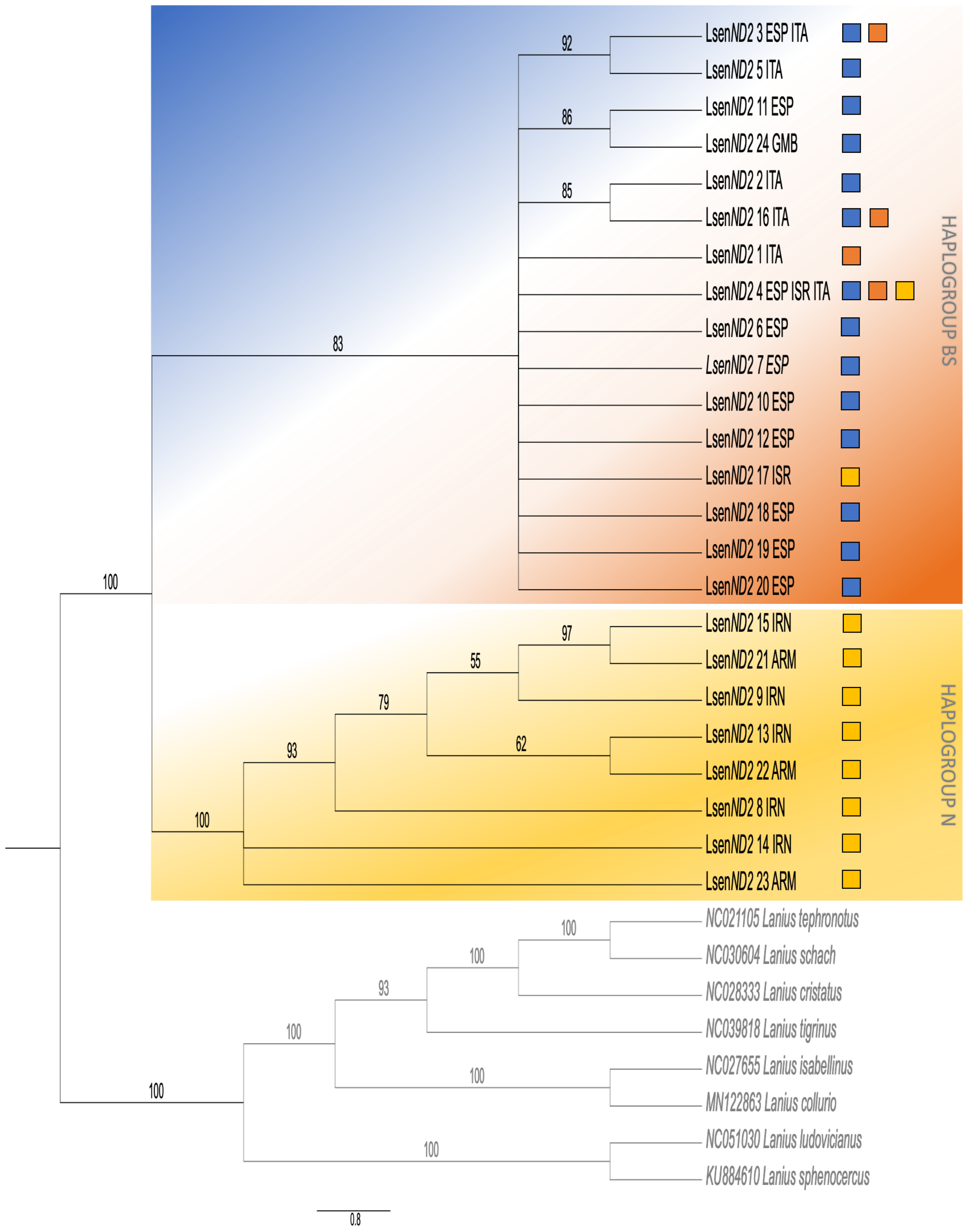

LsenND2 3 ESP ITA
LsenND2 5 ITA
LsenND2 11 ESP
LsenND2 24 GMB
LsenND2 2 ITA
LsenND2 16 ITA
LsenND2 1 ITA
LsenND2 4 ESP ISR ITA
LsenND2 6 ESP
LsenND2 7 ESP
LsenND2 10 ESP
LsenND2 12 ESP
LsenND2 17 ISR
LsenND2 18 ESP
LsenND2 19 ESP
LsenND2 20 ESP
LsenND2 15 IRN
LsenND2 21 ARM
LsenND2 9 IRN
LsenND2 13 IRN
LsenND2 22 ARM
LsenND2 8 IRN
LsenND2 14 IRN
LsenND2 23 ARM
NC021105 Lanius tephronotus
NC030604 Lanius schach
NC028333 Lanius cristatus
NC039818 Lanius tigrinus
NC027655 Lanius isabellinus
MN122863 Lanius collurio
NC051030 Lanius ludovicianus
KU884610 Lanius sphenocercus
HAPLOGROUP BS
HAPLOGROUP N
92
86
85
83
100
97
55
79
62
93
100
100
100
93
100
100
100
100
0.8

S2 (b)

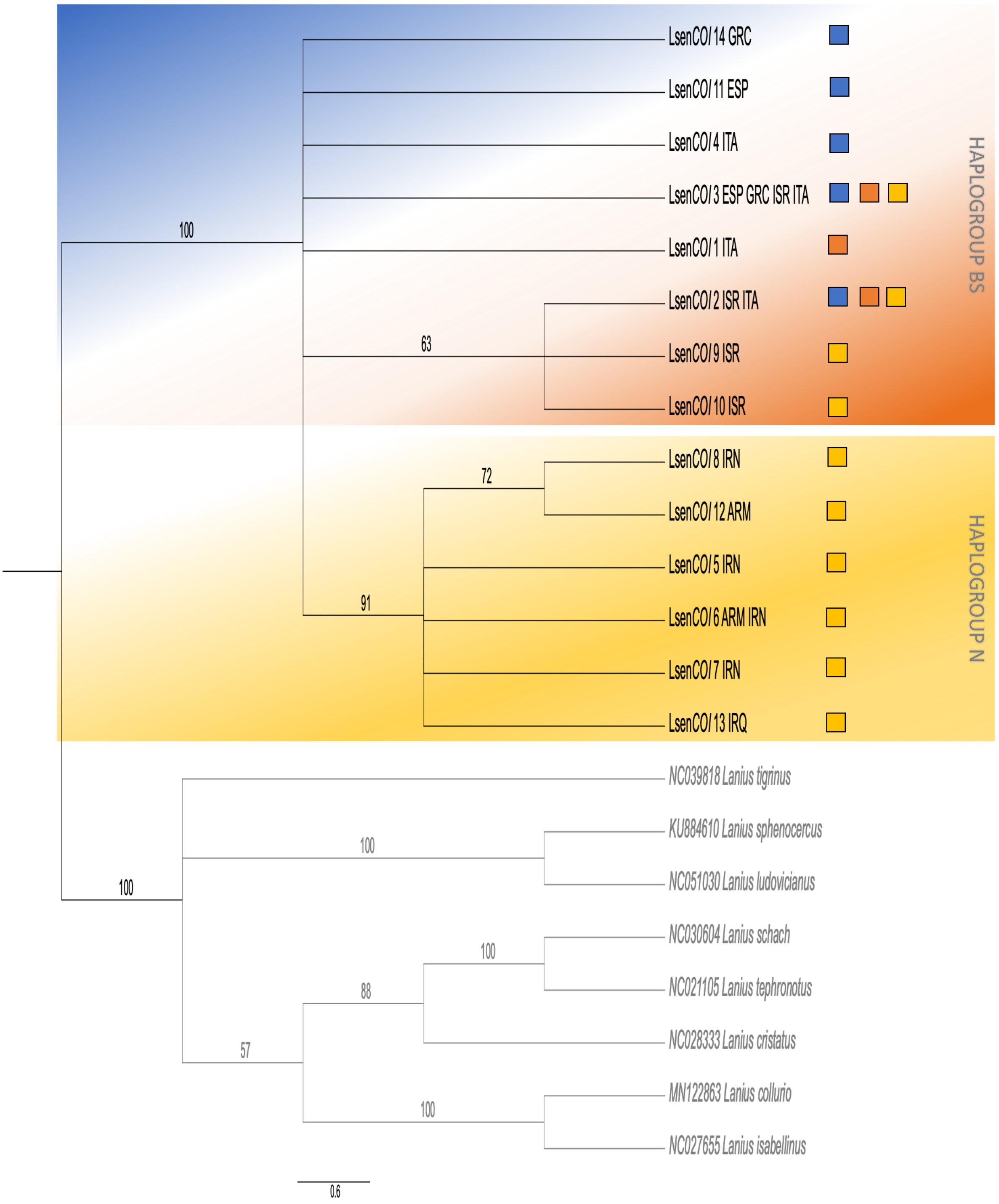
LsenCOI 14 GRC
LsenCOI 11 ESP
LsenCOI 4 ITA
LsenCOI 3 ESP GRC ISR ITA
LsenCOI 1 ITA
LsenCOI 2 ISR ITA
LsenCOI 9 ISR
LsenCOI 10 ISR
LsenCOI 8 IRN
LsenCOI 12 ARM
LsenCOI 5 IRN
LsenCOI 6 ARM IRN
LsenCOI 7 IRN
LsenCOI 13 IRQ
NC039818 Lanius tigrinus
KU884610 Lanius sphenocercus
NC051030 Lanius ludovicianus
NC030604 Lanius schach
NC021105 Lanius tephronotus
NC028333 Lanius cristatus
MN122863 Lanius collurio
NC027655 Lanius isabellinus
HAPLOGROUP BS
HAPLOGROUP N
100
63
72
91
100
100
100
88
57
100
0.6

**S2 (c)**

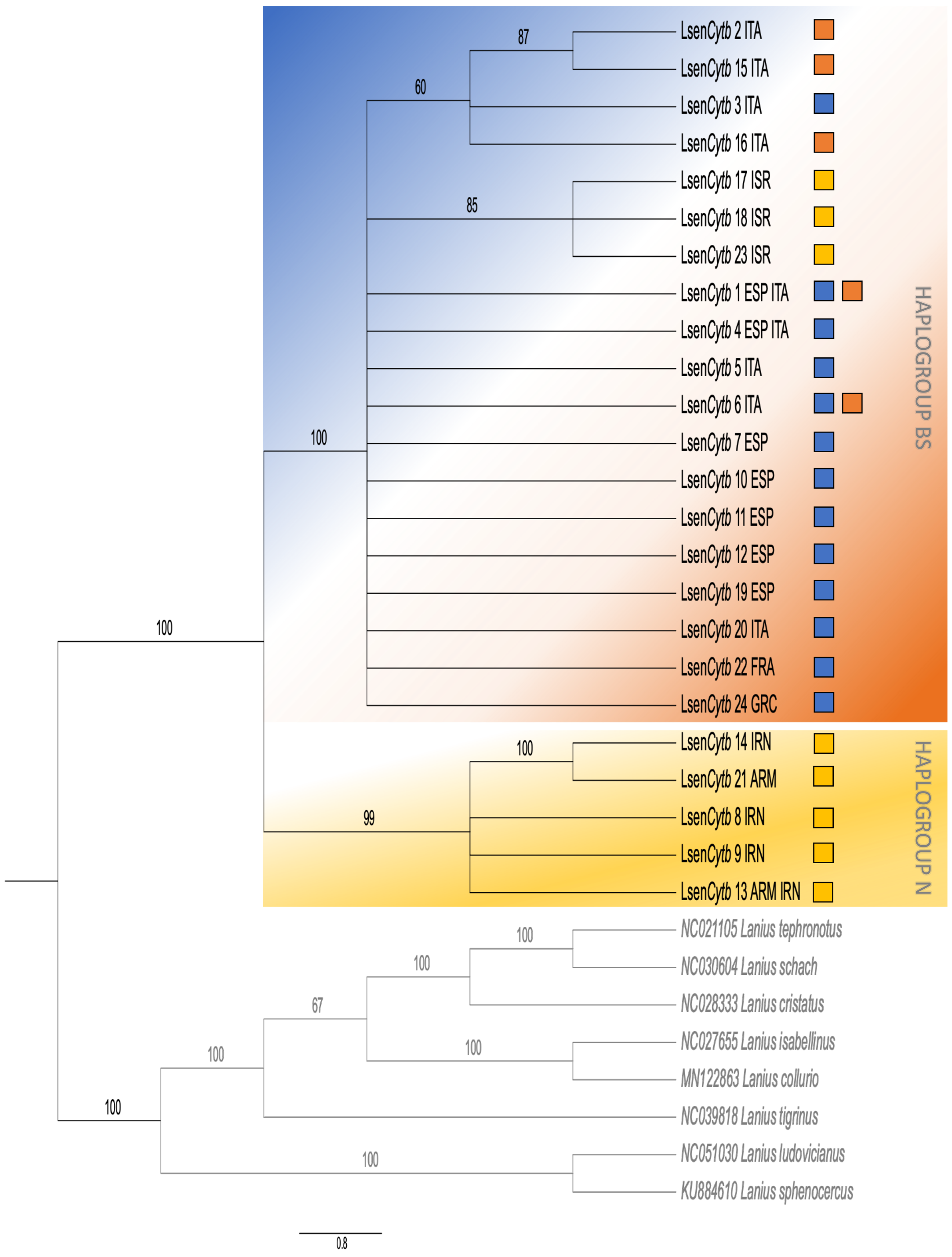
87
60
85
100
100
99
100
100
67
100
100
100
100
100
LsenCytb 2 ITA
LsenCytb 15 ITA
LsenCytb 3 ITA
LsenCytb 16 ITA
LsenCytb 17 ISR
LsenCytb 18 ISR
LsenCytb 23 ISR
LsenCytb 1 ESP ITA
LsenCytb 4 ESP ITA
LsenCytb 5 ITA
LsenCytb 6 ITA
LsenCytb 7 ESP
LsenCytb 10 ESP
LsenCytb 11 ESP
LsenCytb 12 ESP
LsenCytb 19 ESP
LsenCytb 20 ITA
LsenCytb 22 FRA
LsenCytb 24 GRC
LsenCytb 14 IRN
LsenCytb 21 ARM
LsenCytb 8 IRN
LsenCytb 9 IRN
LsenCytb 13 ARM IRN
NC021105 Lanius tephronotus
NC030604 Lanius schach
NC028333 Lanius cristatus
NC027655 Lanius isabellinus
MN122863 Lanius collurio
NC039818 Lanius tigrinus
NC051030 Lanius ludovicianus
KU884610 Lanius sphenocercus
HAPLOGROUP BS
HAPLOGROUP N
0.8

# S2 (d)

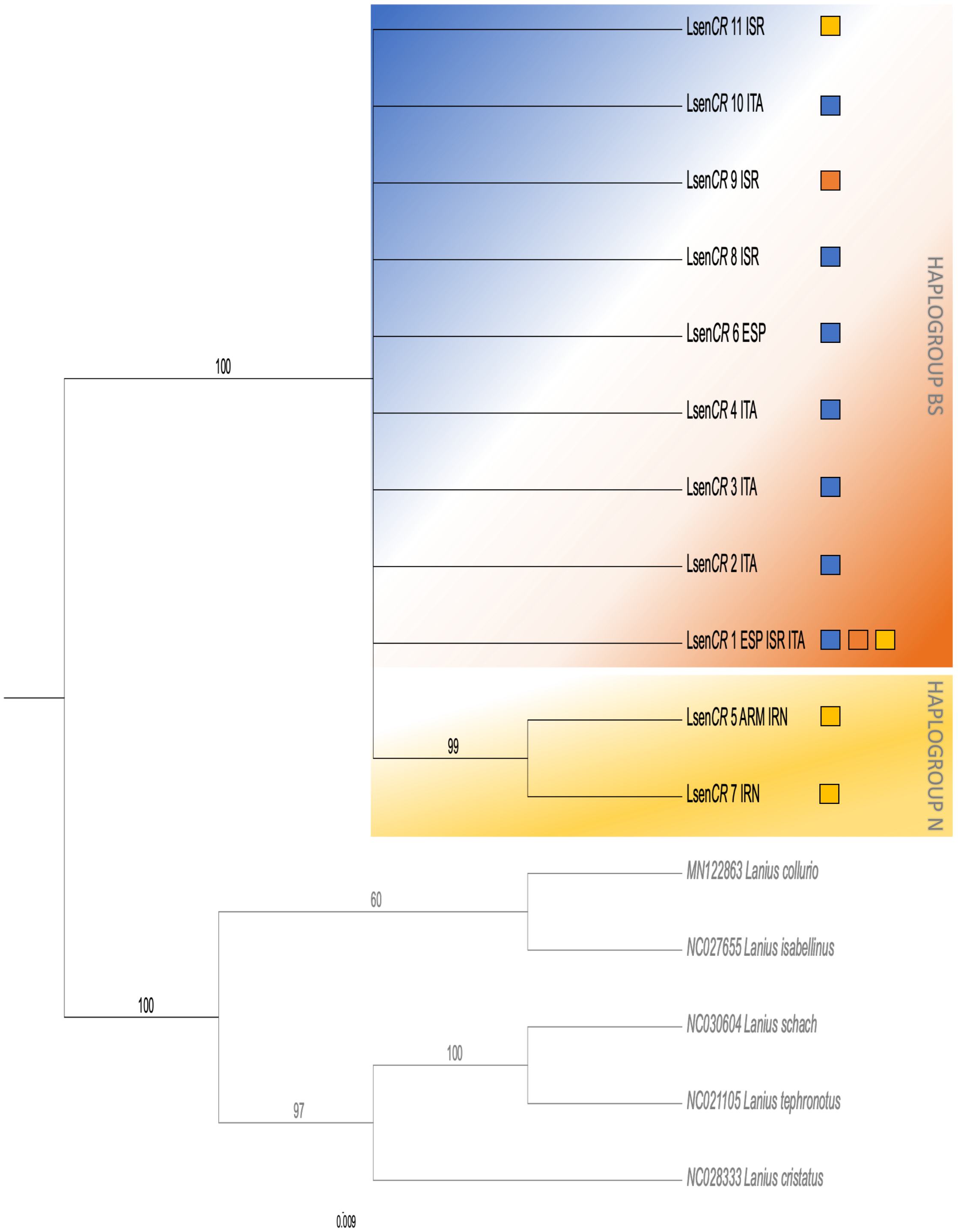
LsenCR 11 ISR
LsenCR 10 ITA
LsenCR 9 ISR
LsenCR 8 ISR
LsenCR 6 ESP
LsenCR 4 ITA
LsenCR 3 ITA
LsenCR 2 ITA
LsenCR 1 ESP ISR ITA
LsenCR 5 ARM IRN
LsenCR 7 IRN
HAPLOGROUP BS
HAPLOGROUP N
MN122863 Lanius collurio
NC027655 Lanius isabellinus
NC030604 Lanius schach
NC021105 Lanius tephronotus
NC028333 Lanius cristatus
100
99
60
100
100
97
0.009

**S3 (a)**

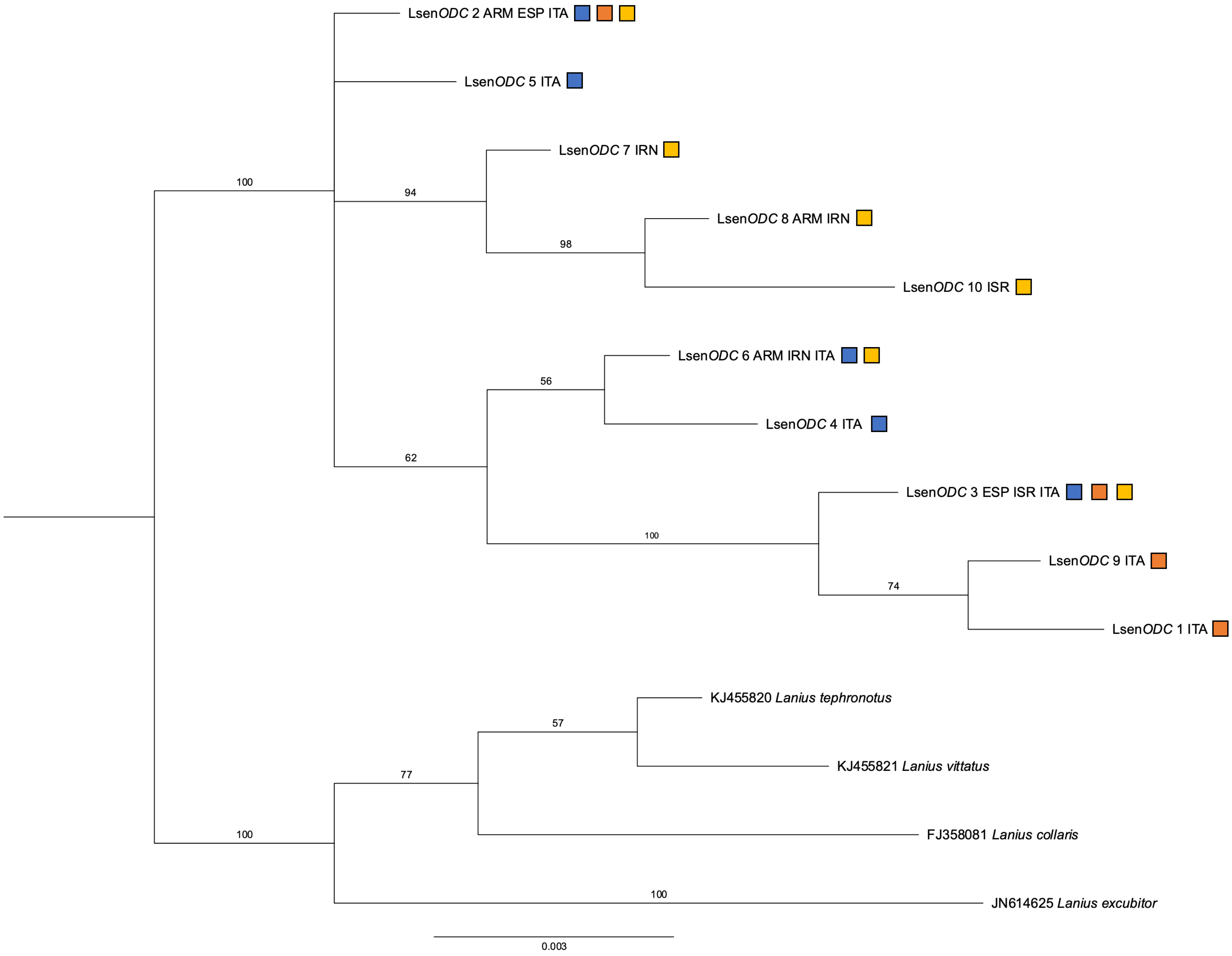
LsenODC 2 ARM ESP ITA
LsenODC 5 ITA
LsenODC 7 IRN
LsenODC 8 ARM IRN
LsenODC 10 ISR
LsenODC 6 ARM IRN ITA
LsenODC 4 ITA
LsenODC 3 ESP ISR ITA
LsenODC 9 ITA
LsenODC 1 ITA
KJ455820 Lanius tephronotus
KJ455821 Lanius vittatus
FJ358081 Lanius collaris
JN614625 Lanius excubitor
100
94
98
56
62
100
74
57
77
100
100
0.003

S3 (b)

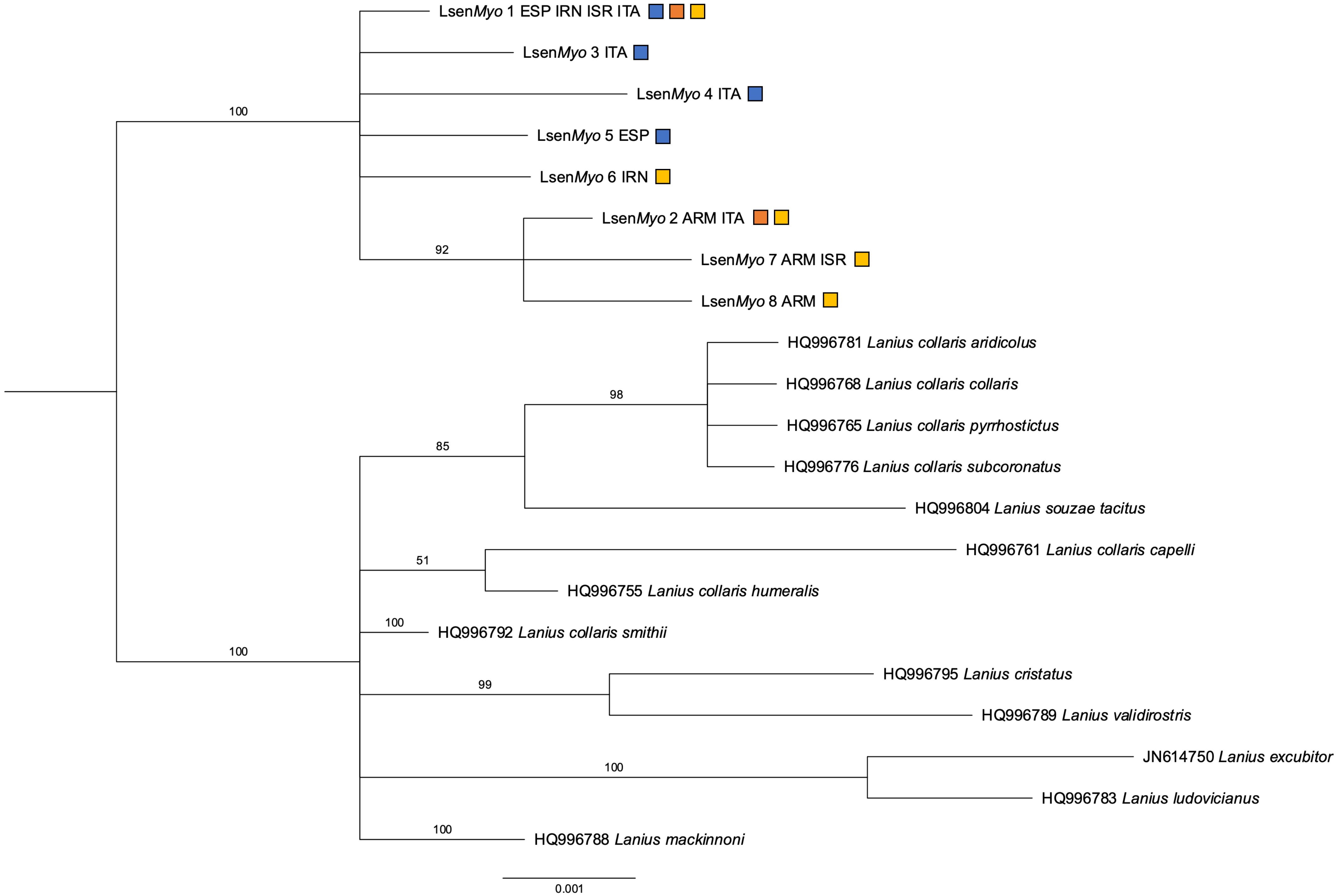
LsenMyo 1 ESP IRN ISR ITA
LsenMyo 3 ITA
LsenMyo 4 ITA
LsenMyo 5 ESP
LsenMyo 6 IRN
LsenMyo 2 ARM ITA
LsenMyo 7 ARM ISR
LsenMyo 8 ARM
HQ996781 Lanius collaris aridicolus
HQ996768 Lanius collaris collaris
HQ996765 Lanius collaris pyrrhostictus
HQ996776 Lanius collaris subcoronatus
HQ996804 Lanius souzae tacitus
HQ996761 Lanius collaris capelli
HQ996755 Lanius collaris humeralis
HQ996792 Lanius collaris smithii
HQ996795 Lanius cristatus
HQ996789 Lanius validirostris
JN614750 Lanius excubitor
HQ996783 Lanius ludovicianus
HQ996788 Lanius mackinnoni
100
92
98
85
51
100
100
99
100
100
0.001

**S4**

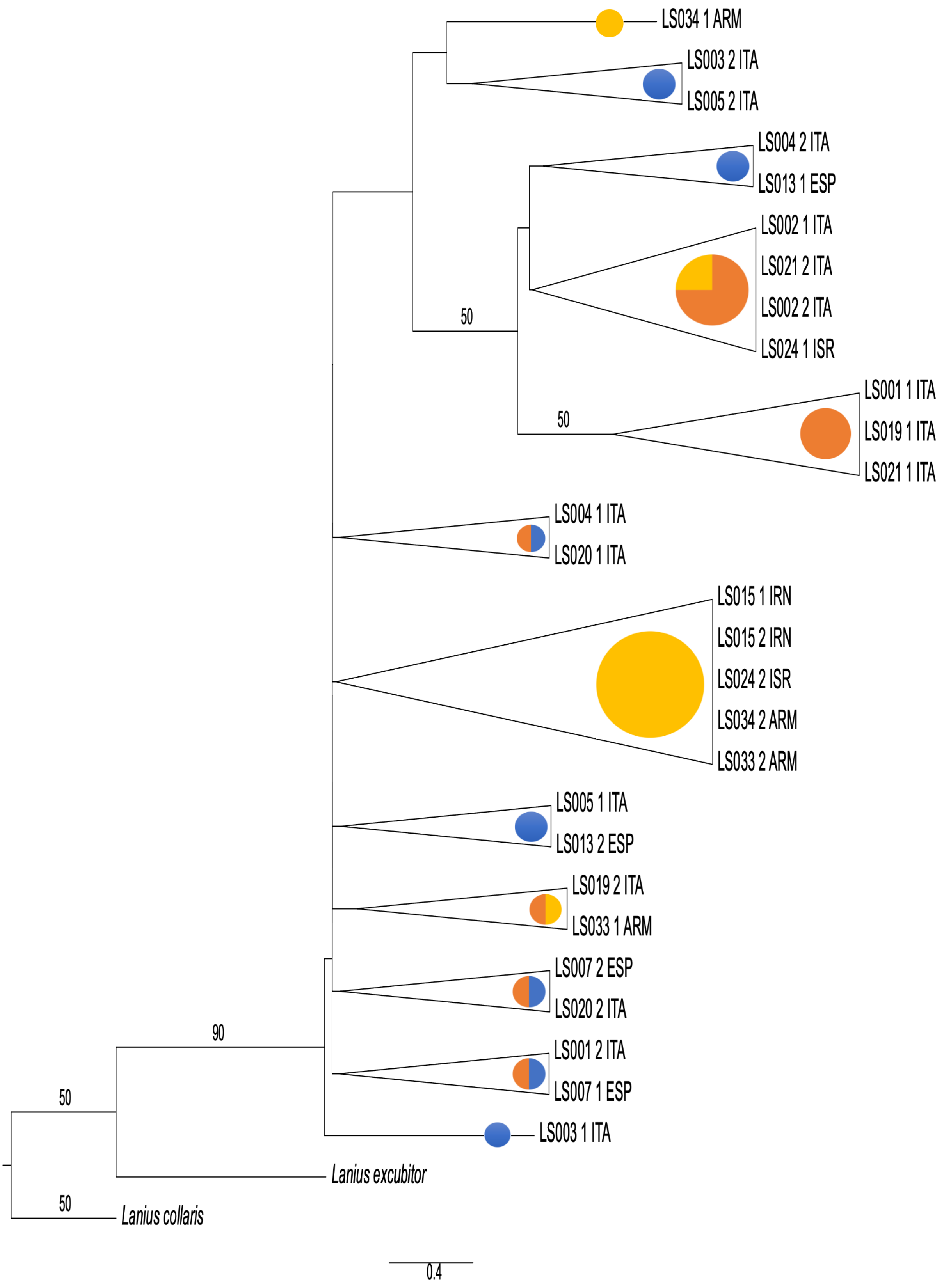

LS034 1 ARM
LS003 2 ITA
LS005 2 ITA
LS004 2 ITA
LS013 1 ESP
LS002 1 ITA
LS021 2 ITA
LS002 2 ITA
LS024 1 ISR
50
LS001 1 ITA
LS019 1 ITA
LS021 1 ITA
50
LS004 1 ITA
LS020 1 ITA
LS015 1 IRN
LS015 2 IRN
LS024 2 ISR
LS034 2 ARM
LS033 2 ARM
LS005 1 ITA
LS013 2 ESP
LS019 2 ITA
LS033 1 ARM
LS007 2 ESP
LS020 2 ITA
LS001 2 ITA
LS007 1 ESP
90
LS003 1 ITA
50
Lanius excubitor
50
Lanius collaris
0.4
